# PathScore: a web tool for identifying altered pathways in cancer data

**DOI:** 10.1101/067090

**Authors:** Stephen G. Gaffney, Jeffrey P. Townsend

## Abstract

**Summary:** *PathScore* quantifies the level of enrichment of somatic mutations within curated pathways, applying a novel approach that identifies pathways enriched across patients. The application provides several user-friendly, interactive graphic interfaces for data exploration, including tools for comparing pathway effect sizes, significance, gene-set overlap and enrichment differences between projects.

**Availability and Implementation:** Web application available at pathscore.publichealth.yale.edu. Site implemented in Python and MySQL, with all major browsers supported. Source code available at github.com/sggaffney/pathscore with a GPLv3 license.

**Supplementary Information:** Additional documentation can be found at http://pathscore.publichealth.yale.edu/faq.

## 1 INTRODUCTION

We present an algorithm and web application, *PathScore*, for the identification of known pathways that are enriched for mutations within a multi-patient somatic gene variant dataset. The algorithm belongs to a class of pathway analysis techniques known as overrepresentation analysis (ORA) tools (Khatri *et al*., 2012). Like other ORA tools, this new algorithm uses a hypergeometric test to estimate pathway alteration probability, but distinguishes itself in three ways. First, it segregates data by patient, calculating patient-specific pathway alteration probabilities that account for varied total mutation counts per patient. Second, it accounts for gene transcript length, incorporating the increased chance of mutation in longer genes. Third, it uses empirically-derived background mutation rates to account for varied mutation probability across the genome, which to our knowledge is a unique feature among pathway analysis tools.

Our web app implementation of the algorithm uses the collection of ‘canonical pathways’ from the Molecular Signatures Database (*MSigDB*, Liberzon *et al*., 2011), which includes pathways from the KEGG, Biocarta, Reactome and Nature-NCI databases, among others. Unlike tools such as MEMo (Ciriello *et al*., 2012) and HotNet2 (Leiserson *et al*., 2014), that detect significantly altered subnetworks, PathScore calculates an enrichment score for every pathway, and all activity in a pathway contributes to the score. In contrast to tools that, for tractability, require stringent gene filtering (MEMo’s filters include a threshold for a minimum patient frequency), PathScore has no filtering requirement. Even a gene that is mutated only once in a dataset will increase the enrichment score if mutated in a patient with no other pathway alterations.

Applications of PathScore to lung adenocarcinoma and lung squamous cell carcinoma data from The Cancer Genome Atlas (Kandoth, 2014) demonstrate its potential to uncover patterns in low frequency events (Fig. 1). Numerous pathways known to be frequently affected in these cancers are identified by PathScore. Additional pathways identified include the nicotinic acetylcholine receptor (nAChR) and GABA receptor pathways (21% and 33% patient coverage, respectively). Hotnet2 results include their genes, but scattered in separate low coverage subnetworks (max 10% and 17% coverage). Both pathways are missed by MEMo due to the low frequencies of their genes. These two related gene families are thought to play a role in lung cancer and have been suggested as potential therapeutic targets (Schuller, 2009). Our demonstration of pathway enrichment supports this hypothesis.

**Fig. 1.**
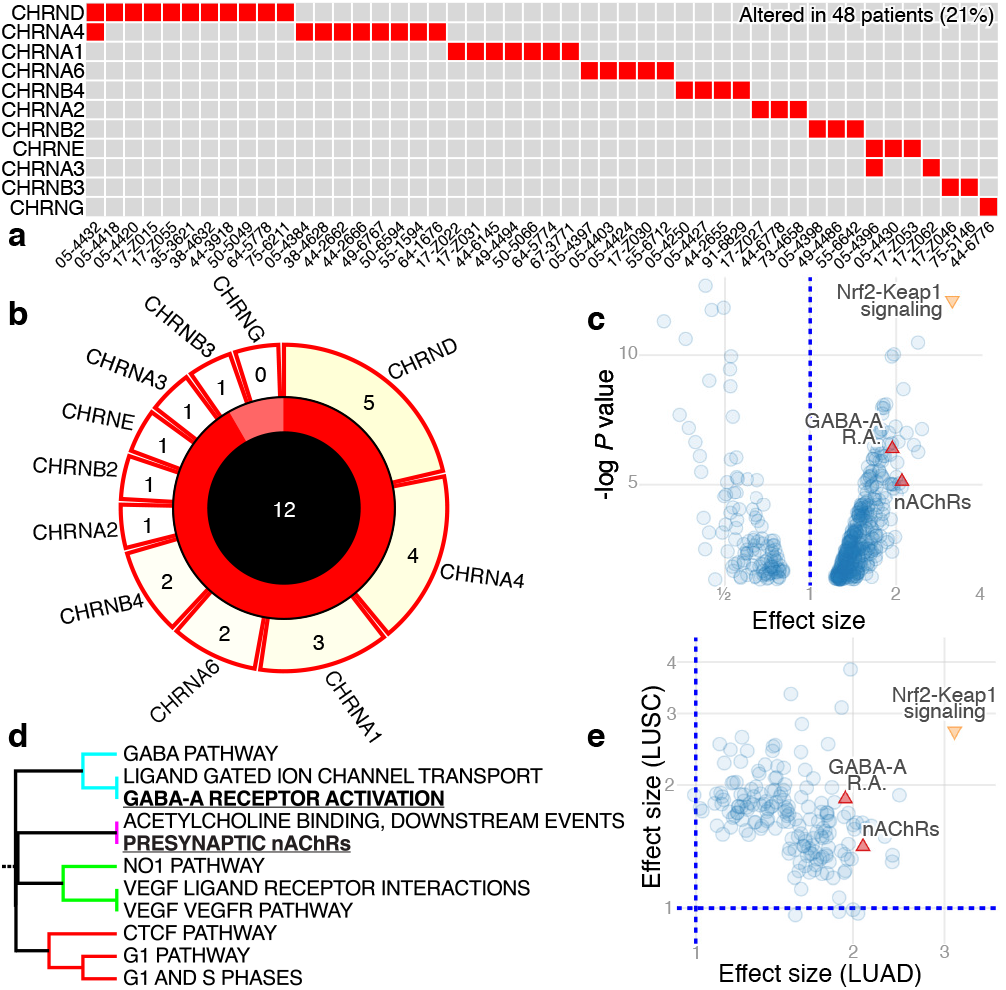
(a) Matrix plot of patient-gene pairs for Reactome’s *Presynaptic nAChRs* pathway in lung adenocarcinoma. (b) Target plot for the same pathway, with a pathway effect size (overburden of mutations) of 2.1. There are 12 genes in the pathway, with 11 mutated in at least one subject, as indicated by the shading of ^11^/_12_ of the pie chart. These genes are named in the tabs whose size and color (via heatmap) are proportional to their respective patient mutation frequency. (c) Volcano plot of altered pathways in lung adenocarcinoma, highlighting the well-known *Nrf2-Keap1 signaling* pathway and two pathways with putative links to cancer: *GABA-A receptor activation* and *Presynaptic nAChRs*. (d) Tree plot detail showing overlap relationships in a subset of lung adenocarcinoma pathways. (e) Comparison plot for lung adenocarcinoma (LUAD) and lung squamous cell carcinoma (LUSC), showing pathways with Bonferroni-adjusted *P* < 0.1.

## 2 APPROACH

### 2.1 The PathScore algorithm

The PathScore algorithm takes as input a set of patient-gene pairs, that should represent all observed non-silent and potentially function-altering mutations. Any listed genes that are not present in the pathways database are discarded.

The algorithm interprets mutations as samples from sites in the genome without explicitly modeling differences in nucleotide sequence context or the functional impact of each mutation. It uses the hypergeometric distribution to calculate the probability that from *M* samples, corresponding to a patient’s *M* mutations, at least one occurs at a site belonging to a particular pathway. We define the number of sites occupied by a set of genes Γ as:

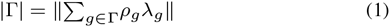

where *ρ_g_* is a gene-specific background mutation rate (BMR), in mutations per megabase, and *λ_g_* is canonical transcript base-length. Scaling by BMR captures the large variation in mutation rate across the genome. As these rates are concordant across tissue types, we use by default an average rate across 91 cancer cell lines (Lawrence *et al*., 2013, Table S5), but allow users to substitute custom BMR values (e.g. tissue-specific rates; Polak *et al*., 2015). For scenarios in which BMR-scaling is inappropriate (e.g. pathway analysis of chromatin marks), *ρ_g_* can be set to 1. For scenarios in which mutation probability doesn’t scale with base-length (e.g. large-scale chromosomal events), genes can be assigned equal mutation probability by setting *λ_g_* to 1.

The formula for gene set size (Eq. 1) can be used to calculate both the target size of a pathway and the background size (*G*) for the set of all genes in the pathways database. For each pathway, PathScore calculates likelihood as the product of patient-specific probabilities, across all *n* patients. Expressed as a function of pathway size, *N*:

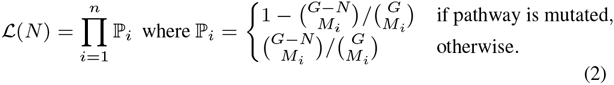

In the null model, *N* is given by the actual pathway size (*N_α_*). PathScore then constructs an alternative model by treating *N* as a free parameter, calculating the maximum likelihood estimate of pathway size (*N*\*), or ‘effective pathway size’. 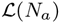 and 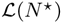 are compared using the likelihood ratio test. The test statistic is assessed by comparison to a chisquare distribution with one degree of freedom, yielding a P value for the pathway—a measure of statistical significance of the disparity between actual and effective pathway sizes. The ratio of effective pathway size to actual pathway size provides a measure of the magnitude of the effect, and is used to rank the relative degree of overburden of pathways with mutation. P values, Bonferroni-adjusted P values, and effect sizes are calculated for each pathway in the database.

### 2.2 The web app

The PathScore app provides both an interactive web interface and a REST API. Users can create a project by submitting a dataset of patient–gene pairs to the ‘Upload’ page. Optional registration entitles users to increased project storage as well as email notification when results are available. Algorithm options are ‘BMR-scaled gene length’ (with gene-specific *ρ_g_* and *λ_g_*), ‘unscaled gene length’ (specifying *ρ_g_* = 1), and ‘gene count’ (specifying *ρ_g_* = 1 and *λ_g_* = 1). Analysis can be customized with two user-specified gene lists: genes to suppress in the analysis, or genes to highlight (i.e. filter out pathways that exclude them). Additional features of the interface include:

#### Pathway visualization

We represent each enriched pathway with two plots. A matrix plot (Fig. 1a) conveys patient-gene pairings and reveals patterns of mutual exclusivity and co-occurrence. Squares in the matrix will exhibit user annotations of gene or mutation attributes in each patient. To convey pathway enrichment attributes, we devised an information-rich, easy to scan ‘target plot’ (Fig. 1b). The target is composed of a black circle with area proportional to the actual pathway size centered within a red circle with area proportional to effective pathway size. The ratio of these areas is the effect size. The plot also indicates the number of genes in the pathway, the mutated gene names, the fraction of the pathway altered, and the percentage of patients with a mutation in each gene.

#### Status page

An overview page with links to browse and download results.

#### Basic results page

All enriched pathway results can be browsed in descending order of effect size.

#### Volcano plot view

Effect sizes and *P* values for all enriched pathways are plotted in an interactive scatter plot (Fig. 1c). Selecting data points reveals pathway information.

#### Tree view

The gene sets of enriched pathways can overlap with each other. Overlapping pathways could be biologically distinct, or they might reflect redundancy in collated pathway databases. Using overlap percentages between all pairs of top pathways, we create an ‘average linkage’ hierarchical cluster tree (Fig. 1d) to summarize distinct patterns in the results and to provide an alternative means of browsing pathways. In the tree view, hovering over leaf nodes displays enrichment information.

#### Project comparison

Users can compare any two uploaded projects (Fig. 1e). Pathways that are enriched in either project are positioned on a log-log scatter plot according to their effect sizes in the two projects. Quadrants around the origin (1,1) distinguish pathways that are enriched in both projects (upper right quadrant) or in only one of the two projects (upper left or lower right quadrant). This plot reveals similarities and differences between cancer types or patient subgroups.

## Acknowledgement

We thank Drs. Benjamin Turk, David Calderwood, Peng Yue, Alessandro Santin, Richard Lifton, and Joseph Schlessinger for their feedback.

